# Ilomastat contributes to the survival of mouse after irradiation via promoting the recovery of hematopoietic system

**DOI:** 10.1101/2020.08.13.249235

**Authors:** Baoquan Zhao, Xiaoman Li, Xingzhou Li, Dongqin Quan, Fang Zhang, Burong Hu

## Abstract

Ilomastat, a broad-spectrum inhibitor of matrix metalloproteinases (MMPs), has drawn attentions for its function in alleviating radiation damage. However, the detailed mechanisms of Ilomastat’s protection from animal model remain not fully clear. In this study, the C57BL/6 mice were pre-administrated with Ilomastat or vihicle for 2 h, and then total body of mice were exposed to 6 Gy of γ-rays. The protective effect of Ilomastat on the hematopoietic system in the irradiated mice were investigated. We found that pretreatment with Ilomastat significantly reduced the level of TGF-β1 and TNF-α, and elevated the number of bone marrow (BM) mononuclear cells in the irradiated mice. Ilomastat pretreatment also increased the fraction of BM hematopoietic progenitor cells (HPCs) and hematopoietic stem cells (HSCs) at day 30 after irradiation, and protected the spleen of mouse from irradiation. These results suggest that Ilomastat promotes the recovery of hematopoietic injury in the irradiated mice, and thus contributes to the survival of mouse after irradiation.

## 1 Introduction

Our previous studies showed that Ilomastat, a broad-spectrum inhibitor of matrix metalloproteinases (MMPs), could improve the survival of mice after γ-ray irradiation. However, the detailed mechanisms underlying the protective effect of Ilomastat on animal model remain largely unknown. High-dose ionizing irradiation (IR) can result in severe acute radiation syndrome (ARS) [1–3]. Depletion of peripheral blood cells and immunosuppression are the main characters of ARS [4, 5]. A number of compounds have been reported to reduce radiation-induced hematopoietic injury and improve the survival of the irradiated mice [4, 6]. The extracellular matrix (ECM), a major component of hematopoietic microenvironment, which alternates in the structure and/or function may contribute to the development of hematologic disorders [7–9]. MMPs’ functions have been researched widely, which mainly focus on inflammation and cancer progression [10, 11]. While MMPs also can modulate the proliferation, differentiation, and migration of hematopoietic stem/progenitor cells of different lineages through ECM proteolysis and growth factors/cytokines release [12]. In addition, MMP9 can proteolytically cleave latent TGF-β1 (transforming growth factor-β1) or pro-TNF-α (tumor necrosis factor-α) providing an alternative pathway for activation of TGF-β and TNF-α [13, 14]. TGF-β1 can induce apoptosis of normal progenitors [15]. Chronic injection of TNF-α into animals results in the rapid development of anaemia [16, 17]. It was also reported that radiation-induced increase of MMPs activity caused endothelial cells apoptosis [18] and bone marrow endothelial cells (BMEC) plays a critical role in the regulation of haemopoiesis [19].

Therefore, administration of Ilomastat, inhibitor of MMPs, might improve the hematopoiesis disorders in the irradiated mice and provide the protective effect. To confirm this speculation, we deeply investigated the changes of biomarkers associated to hematopoiesis in the mice treatmented with/without Ilomastat before 6 Gy of total body γ-ray irradiation. Our findings indicated that Ilomastat administration could protect the hematopoietic and immunological system of mice from irradiation.

## 2 Materials and methods

### 2.1 Mice

Male C57BL/6 mice (SPF class) were purchased from Beijing SPF Bioscience Co. Ltd. All mice were used at approximately 6–8 weeks of age and weighting 20 ± 2 g. The animals were kept in Laboratory Animal Center, the Academy of Military Medical Sciences for one week prior to the initiation of this study. The mice had free access to pellet food and water and were housed at a temperature of 20-24 °C and relative humidity of 55 ± 5% on a 12-hour light/12-hour dark cycle.

Animal welfare and experimental procedures were carried out in accordance to National Institutes of Health guidelines for the care and use of laboratory animals. This study was approved by Beijing Experimental Animal Ethics Committee (2006) No. 5118 set by the Beijing People’s Government. Animals in this study were sacrificed by cervical dislocation. All efforts were made to minimize discomfort, distress, pain and injury.

### 2.2 Preparation and administration of Ilomastat

Ilomastat (Sigma-Aldrich, St. Louis, MO), a broad spectrum MMPs inhibitor, was dissolved in a special solvent made by our institute [20]. For Ilomastat administration, mice were given 10 mg/kg of Ilomastat via intraperitoneal injection 2 h before irradiation referring to our previous study [21].

### 2.3 Irradiation treatment of mice

Our previous study showed that a dose of LD50 at day 30 after γ ray exposure to C57BL/6 mice was 6 Gy [21]. Thus, mice were subjected to 6 Gy for the studies of protection mechanism of Ilomastat against radiation in the current study. Mice were equally divided into three groups at random: sham control (administrated with 10 mg/kg of Ilomastat only), IR (administrated with vehicle and irradiated with 6 Gy γ rays) and Ilomastat (10 mg/kg)+IR (6 Gy) groups. Mice were injected intraperitoneally once either with Ilomastat or vehicle control 2 h before and then subjected to total body irradiation (TBI) in the ^60^Co irradiator (Model GWXJ80, NPIC, Chengdu, China) at a dose rate of 0.91 Gy/min. All irradiations were carried out at room temperature. After irradiation, animals were returned to the animal facility for daily observation and subsequent experiments.

### 2.4 Cytokine assay

Assay of cytokines in serum for TGF-β1, stem cell factor (SCF) and TNF-α were performed with Enzyme-Linked Immunosorbent Assay (ELISA). ELISA kits according to the manufacturer’s protocol. Mouse TGF-β1 and SCF ELISA kits were purchased from Wuhan Boster Biological Technology Co.Ltd (Wuhan, China). Mouse TNF-α ELISA kit was purchased from Beijing Dakewe Biotechnology, Inc (Beijing, China).

### 2.5 Organ coefficient calculation

The spleen were collected and weighted after mice were euthanized. The spleen coefficient were calculated by dividing individual organ weight (mg) with their body weight (g).

### 2.6 Endogenous spleen colony forming unit (e-CFU-S)

The classical e-CFU-S was performed essentially as described previously [22]. Spleens were removed from mice at the 10^th^ day post irradiation and fixed in Bouin’s solution (trinitrophenol and methanal) for 24 h. Macroscopic colonies (CFU-S) visible to the naked eye were scored in each spleen.

### 2.7 Bone marrow nucleated cells (BMNC) counts

Mouse femoral bones were collected and bone marrows were flushed out with Hank’s balanced salt solution containing 3% fetal calf serum. Red blood cells (RBCs) were lysed once for 5 min at 4 °C in 0.15 M NH_4_Cl, washed once with PBS (Gibco). The number of bone marrow nucleated cells was counted using a light microscope.

### 2.8 Analysis of proliferation status of cells in the bone marrow

Mice were euthanized at day 3 and 21 post irradiation. Femurs were isolated and bone marrow cells collected as described above. The cells were fixed in 70% chilled ethanol and cell cycle distribution was analyzed to assess the S-phase cells. Briefly, about 1×10^6^ fixed cells were washed in PBS, and subsequently stained with Propidium Iodide (with RNAase A) in PBS for 15 min. Measurements were made with a laser based (488 nm) flow cytometer (FACS Calibur; Becton and Dickinson, USA) and data acquired using the Cell Quest soſtware (Becton and Dickinson, USA). Cell cycle analysis was performed using the Mod Fit program (Becton and Dickinson, USA).

### 2.9 Analysis of BM lineage-sca1^+^ c-kit^+^ cells (indicator of HSCs) and lineage^−^ sca1^−^ c-kit^+^ cells (indictor of HPCs) by flow cytometry

For staining lineage-committed cells in bone marrow, bone marrow mononuclear cells (BM-MNCs) were incubated with Fc block (BD Pharmingen) for 15 min and then incubated with APC-conjugated antibodies specific for murine CD3 (Clone.145-2C11), Mac1/CD11b (Clone.M1/70), CD45R/B220 (Clone.RA-6B2), TER-119 (Clone.TER-119) and Gr-1 (Clone.RB6-8C5) for 15 min at 4 °C. For the staining of hematopoietic stem/progenitor cells (HSPCs), the cells were incubated with PE-conjugated antibodies to c-Kit (Clone. 2B8), PE-Cy7-conjugated antibodies to Sca1 (Clone. D7) and analyzed by flow cytometry. For calculating compensation, we used the stained BD ComBads. In addition, we recorded data from isotype control cells in order to identify background staining. Meanwhile, we also used 7-amino-actinomycin D (7-AAD), a DNA dye that is used to discriminate dead cells from live cells. The percentage of positive cells was counted in BM-MNCs. For each sample, a minimum of 100,000 BM-MNCs were acquired and the data were analyzed using FlowJo 10.2 software (Tree-Star Inc., Ashland, OR), and results are shown as pseudocolor.

### 2.10 Histological examination

Mice were euthanized at day 10, 20 and 30 post irradiation. Bone and spleen were isolated, and paraffin sections were stained with hematoxylin-eosin according to standard procedure. Femurs were then fixed in 10% neutral-buffered formalin (SRL) followed by decalcification by immersing the femurs in 10% EDTA (pH 7.0 –7.4) at 4 °C for 4-5 d. Megakaryocytes were quantified according to morphologic criteria as counted in 9 random fields, from 2 different sections of a femur, per mouse.

### 2.11 Statistical analysis

The results were expressed as the mean ± standard deviations (SD) from at least three independent mice. The data were analyzed by Student’s t-test or one-way analysis of variance (ANOVA). *P* value of less than 0.05 was considered significant.

## 3 RESULTS

### 3.1 Ilomastat pretreatment reduces the levels of hematopoietic inhibitory factor (HIF) and increases the levels of hematopoietic growth factor (HGF) in serum of the irradiated mice

To indicate the protective effects of Ilomastat on the irradiated mice mightily via improving the hematopoiesis, we first investigated the expression of hematopoiesis-related cytokines in the irradiated mice with or without Ilomastat pretreatment. Radiation treatment alone significantly increased the levels of TGF-β1 and TNF-α at day 10 and 20, and decreased the level of SCF in the serum at day 30, compared to the sham control (Fig. 1). Ilomastat pretreatment significantly reduced the levels of hematopoietic inhibitory factors TGF-β1 and TNF-α at day 10 (Fig. 1A and 1F). On the other hand, Ilomastat significantly elevated the level of SCF (Fig. 1G). These results demonstrate that Ilomastat pretreatment could significantly reduce the levels of HIF and increase the levels of HGF in serum of the irradiated mice.

**Figure 1.**
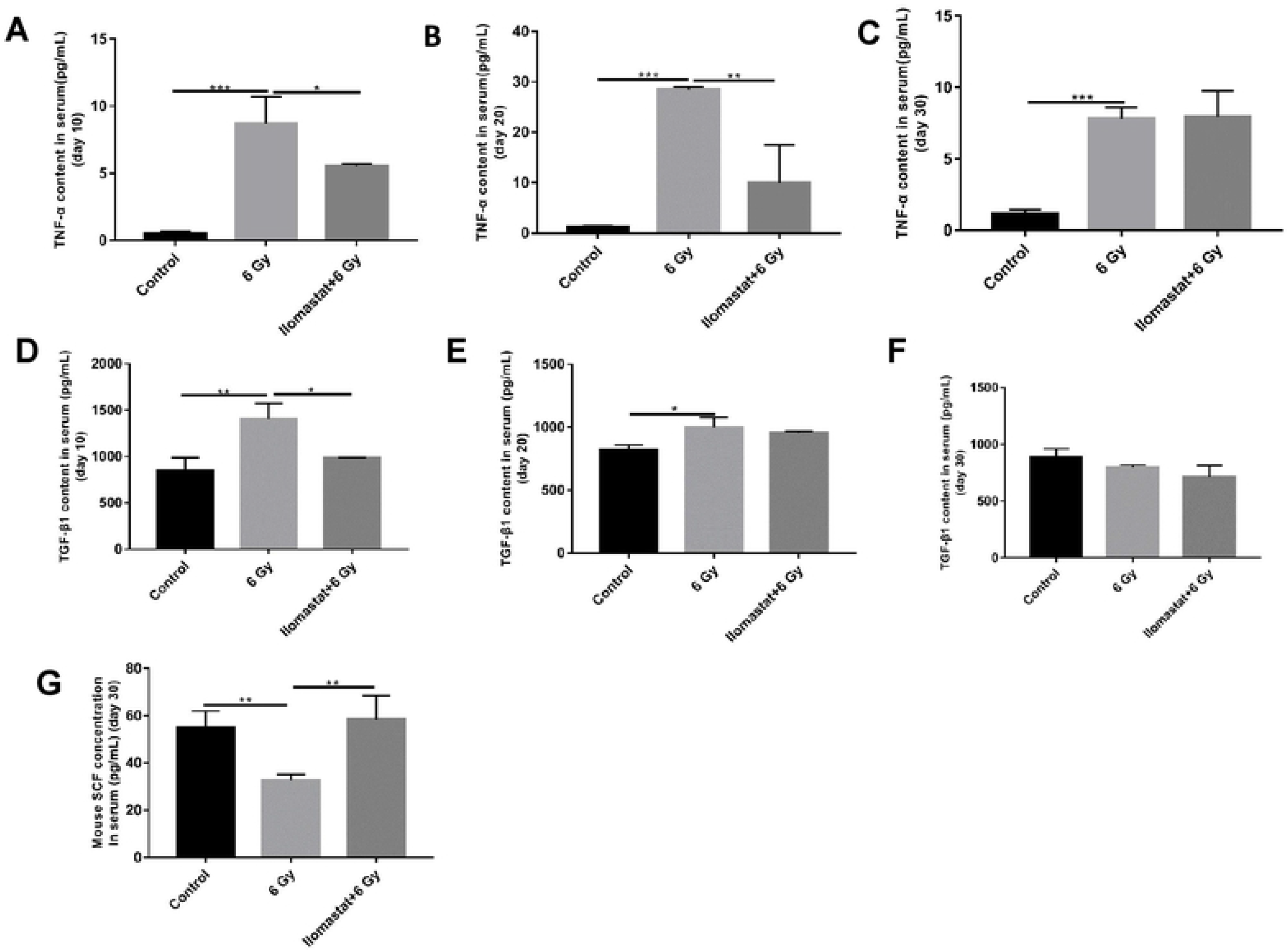
Ilomastat pretreatment influences the cytokines level. TNF–α (**A**, day 10; **B**, day 20; **C**, day 30), TGF-β1 (**D**, day 10; **E**, day 20; **F**, day 30) and SCF (**G**, day 30) levels in mouse serum were measured by colorimetric enzyme-linked immunosorbent assay (ELISA). The final concentration of TGF-β1, TNF-α and SCF in each serum sample were corrected for sample dilution and presented in pg/mL. All error bars indicate SEM (n = 8). * *P* < 0.05; ** *P* < 0.01; *** *P* < 0.001.

### 3.2 Ilomastat pretreatment improves the recovery of bone marrow cellularity

Radiation-induced hematopoietic injury is characterized by loss of the BM cellularity. The effects of Ilomastat on the radiation-induced damage to BM were analyzed by histological examination of the femurs at day 10, 20 and 30 post-TBI. Severe radiation-induced disruption of vasculature and excessive ablation of cellular content of the BM in TBI mice were observed (Fig. 2A). In contrast, Ilomastat pretreatment reversed the radiation-induced cellular depletion to a large extent with an almost similar levels of cell numbers and cellular content in the control mice (Fig. 2A), indicating that Ilomastat pretreatment mitigates the radiation-induced BM ablation of cellularity and accelerates the hematopoietic recovery. We also determined the number of BMNCs at day 10 and 20 post-TBI, and both the BMNC count decreased in the irradiated mice only. On the contrary, the BMNC count in the irradiated mice pretreated with Ilomastat was higher than that in the irradiated mice alone (Fig. 2B and 2C). Furthermore, megakaryocyte counts in bone marrow sections of the irradiated mice plus the Ilomastat pretreatment were significantly higher than that in the sham treated mice (Fig. 2D). Taken together, these results provide an evidence that Ilomastat pretreatment ameliorate the loss of BM cellularity induced by irradiation and increase the proliferative capacity of the BM cells.

**Figure 2.**
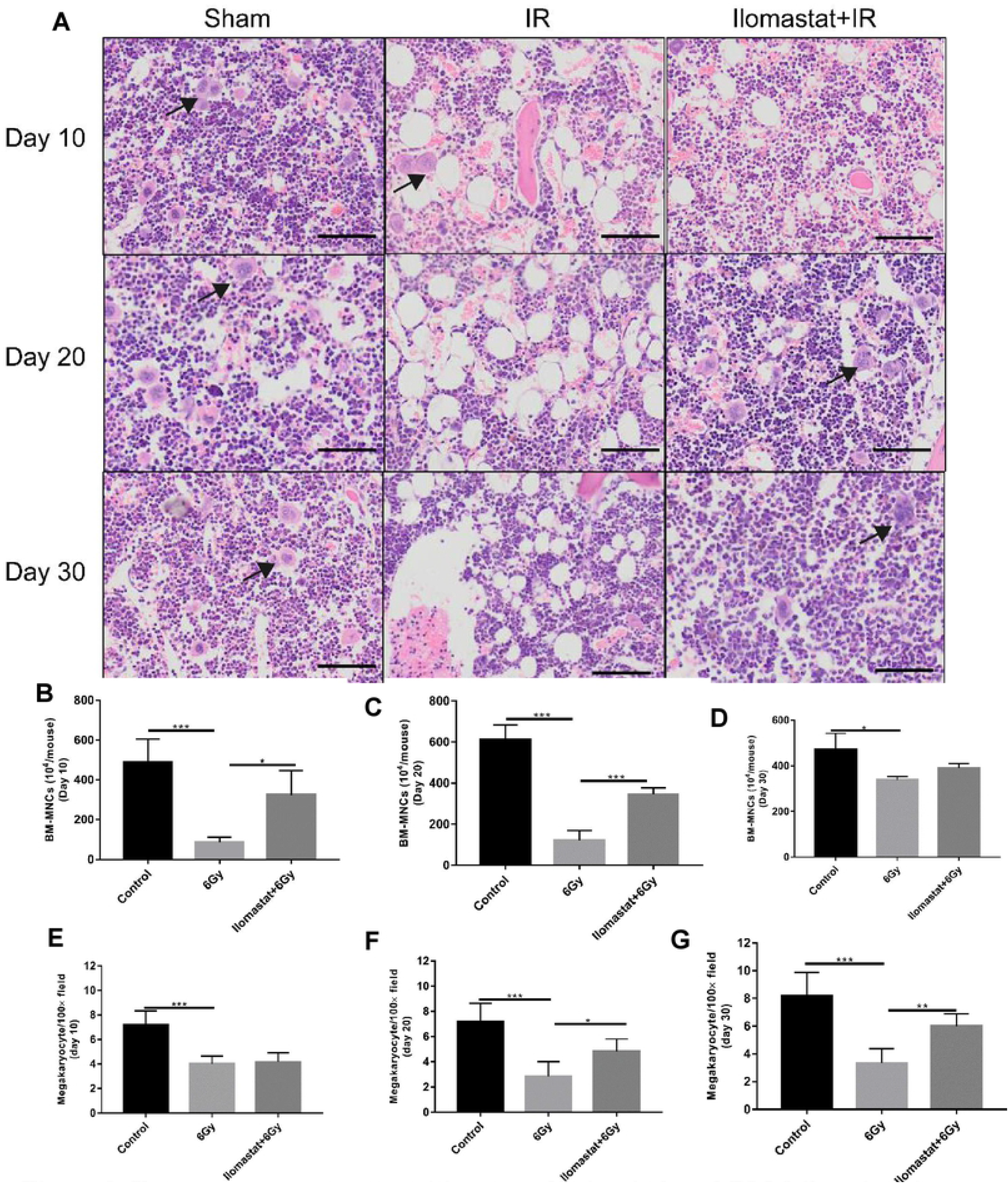
Ilomastat pretreatment mitigates radiation-induced BM failure in mice. The bone marrows were collected from femoral bones of experimental mice and analyzed. **(A)** Panels show H&E staining of mouse femurs at day 10, 20 and 30 after 6 Gy γ-ray irradiation. Black arrowheads point at megakaryocytes. Representative images are shown from control, IR and IR + Ilomastat pretreatments. Total nucleated cell numbers of BM were scored in control, IR and IR + Ilomastat mice at day 10 **(B)** and day 30 **(C)** post TBI under light microscope. **(D)** Number of megakaryocytes in bone sections, determined by counting of 9 random fields per bone section for each mouse. All error bars indicate SEM (n = 8). * *P* < 0.05; ** *P* < 0.01; *** *P* < 0.001.

### 3.3 Ilomastat pretreatment facilitates the proliferations of BM cells

BM stem cells entry into the cell cycle results in hematopoietic recovery. We analyzed the effects of Ilomastat on TBI-induced hematopoietic recovery of BM by analyzing the cell cycle distribution. Ilomastat pretreatment significantly enhanced the fraction of S-phase cell in the irradiated mice at day 3 post-TBI (Fig. 3A/first panel and 3B). The irradiated mice were incapable of combating the radiation-induced damage even at day 20 (lower S-phase cells) as opposed to the mice administered with Ilomastat (Fig. 3A/second panel and 3C). Cumulatively, these results suggest that Ilomastat pretreatment promotes the proliferation of BM cells in the irradiated mice, which appear to serve as potential contributing factors for the protection of irradiation-induced hematopoietic injury.

**Figure 3.**
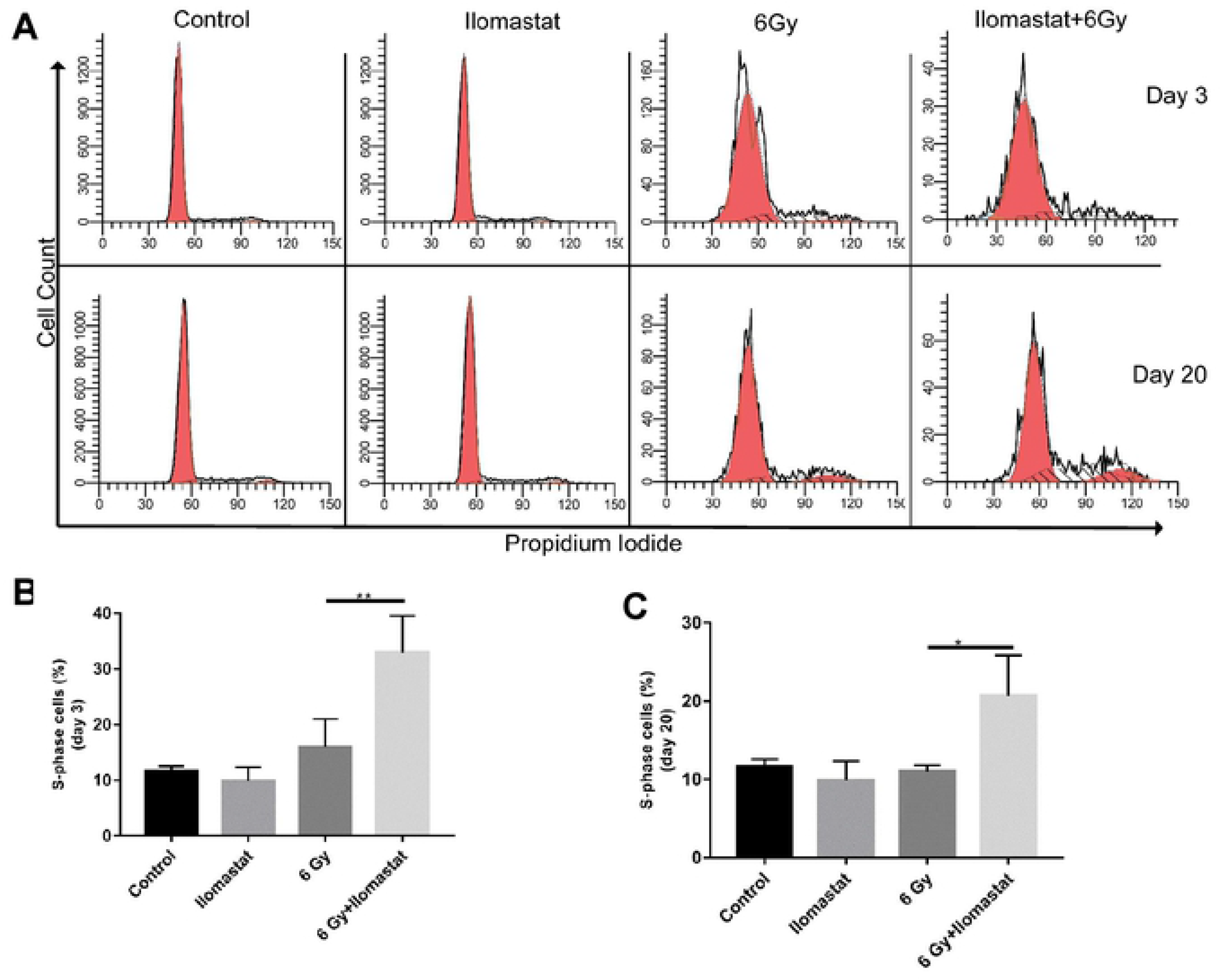
Ilomastat pretreatment stimulates the cell proliferation of BM in irradiated mice. Effect of Ilomastat on radiation mediated compromised proliferative ability in BM was evaluated by analysis of S-phase cells in the BM. Representative DNA flow cytograms **(A)**. Frequency of S-phase cells in the BM of mice at day 3 **(B)** and 20 **(C)** post 6 Gy γ-ray irradiation. \**P* < 0.05, \*\**P* < 0.01. n = 8 in all groups.

### 3.4 Ilomastat promotes hematopoietic stem and progenitor cell recovery after irradiation

It is well established that HSPCs play predominant roles in radioprotection [23]. We assessed the repopulation of HSPCs in bone marrow of mice on day 30 after 6 Gy irradiation. The fractions of hematopoietic stem (HSCs, Lin^−^ Sca-1^+^ c-Kit^+^) and progenitor (HPCs, Lin^−^ Sca-1^−^c-Kit^+^) were determined by flow cytometric assays (Fig. 4A). As shown in figure 4B and 4C, the percentage of HSCs and HPCs slightly decreased by day 30 in the irradiated mice group. While a higher percentage of HSCs and HPCs among the total BM cells was observed in the irradiated mice plus Ilomastat pretreatment group, compared with those in unirradiated control group and irradiated group alone.

**Figure 4.**
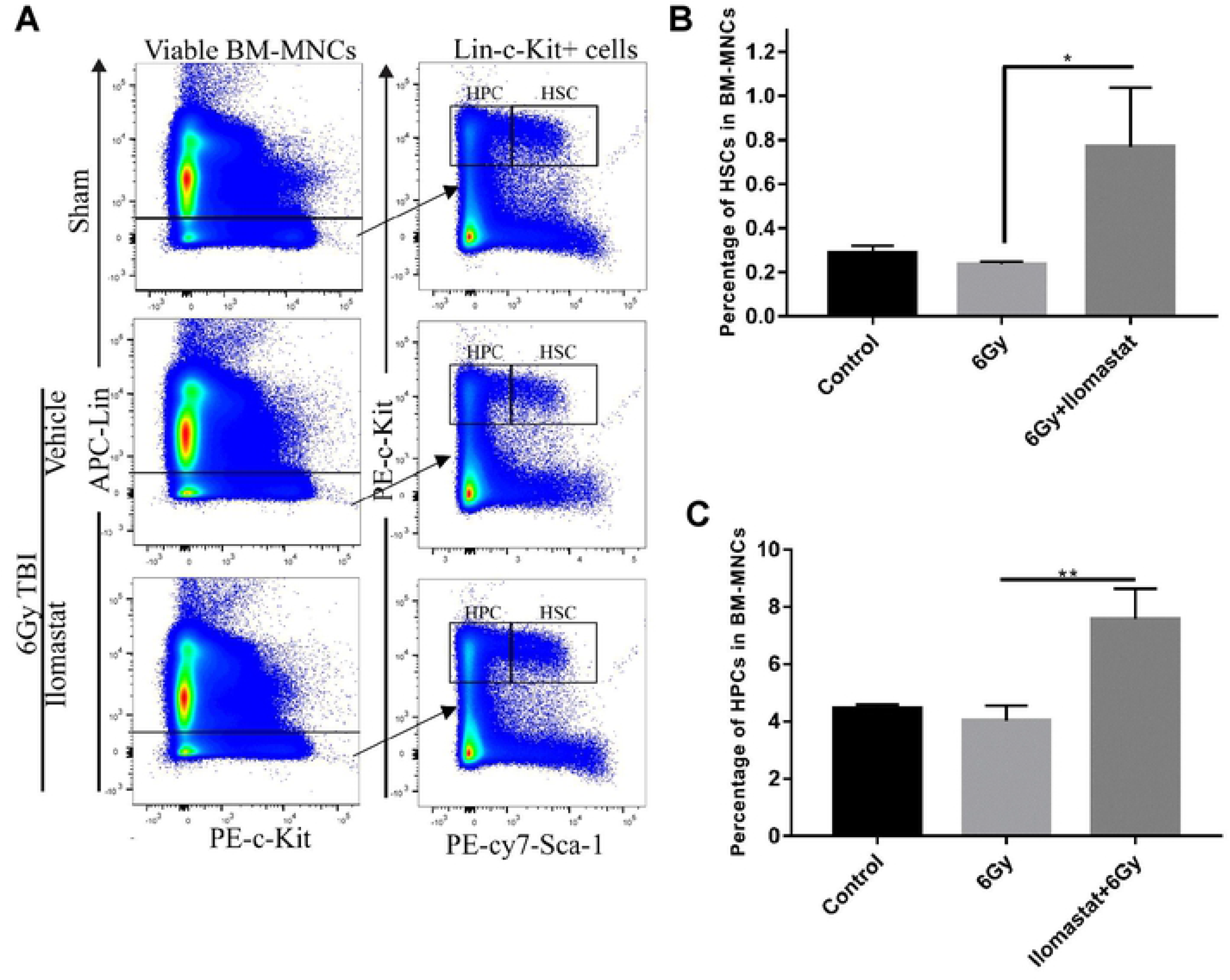
Ilomastat attenuates radiation-induced reduction of HSCs and HPCs. Mice were administrated with/without 10 mg/kg Ilomastat 2 h before exposure to 6 Gy γ-ray irradiation. **(A)** A representative gating strategy of HSC and HPC analysis by flow cytometry. The frequencies of HSCs **(B)** and HPCs **(C)** in BMMCs were presented as mean ± SEM (n = 8). \*\**P* < 0.05.

### 3.5 Ilomastat protects the spleen of irradiated mice and enhances the restoration of hematopoietic potency

It is well known that spleen is an important hematopoietic organ besides bone marrow. There was a clear marginal zone between the white pulp (WP) and red pulp (RP) in control mice (Fig. 5, first column). RP areas dramatically elevated in the spleens of the irradiated mice at day 10 (Fig. 5, second column/first panel). Especially at day 20, the marginal zone between RP and WP were poorly visible (Fig. 5, second column/second panel). At day 30, RP continued to dominate over WP, and a relative increase in WP in the spleens was observed compared to those at day 10 and 20 (Fig. 5, second column/third panel). Compared to the irradiated groups alone at day 20, the damages of the spleen structure in the drug pretreated mice followed by irradiation were dramatically improved (Fig.5, third column/second panel). In addition, the irradiated mice at day 20 exhibited obvious splenomegaly and Ilomastat pretreatment mitigated the radiation-induced splenomegaly (Fig. 6A and 6C). The ratio of spleen to body weight markedly reduced in the irradiated mice at day 10, while it was comparable in the cohort administered Ilomastat following radiation to the sham control mice (Fig. 6E). However, the spleen coefficients of irradiated mice receiving vehicle pretreatment were greater than those from all the other groups at day 20 post 6 Gy radiation (Fig. 6F). These results suggest that Ilomastat can effectively protect another hematopoietic organ (spleen) of the irradiated mice.

**Figure 5.**
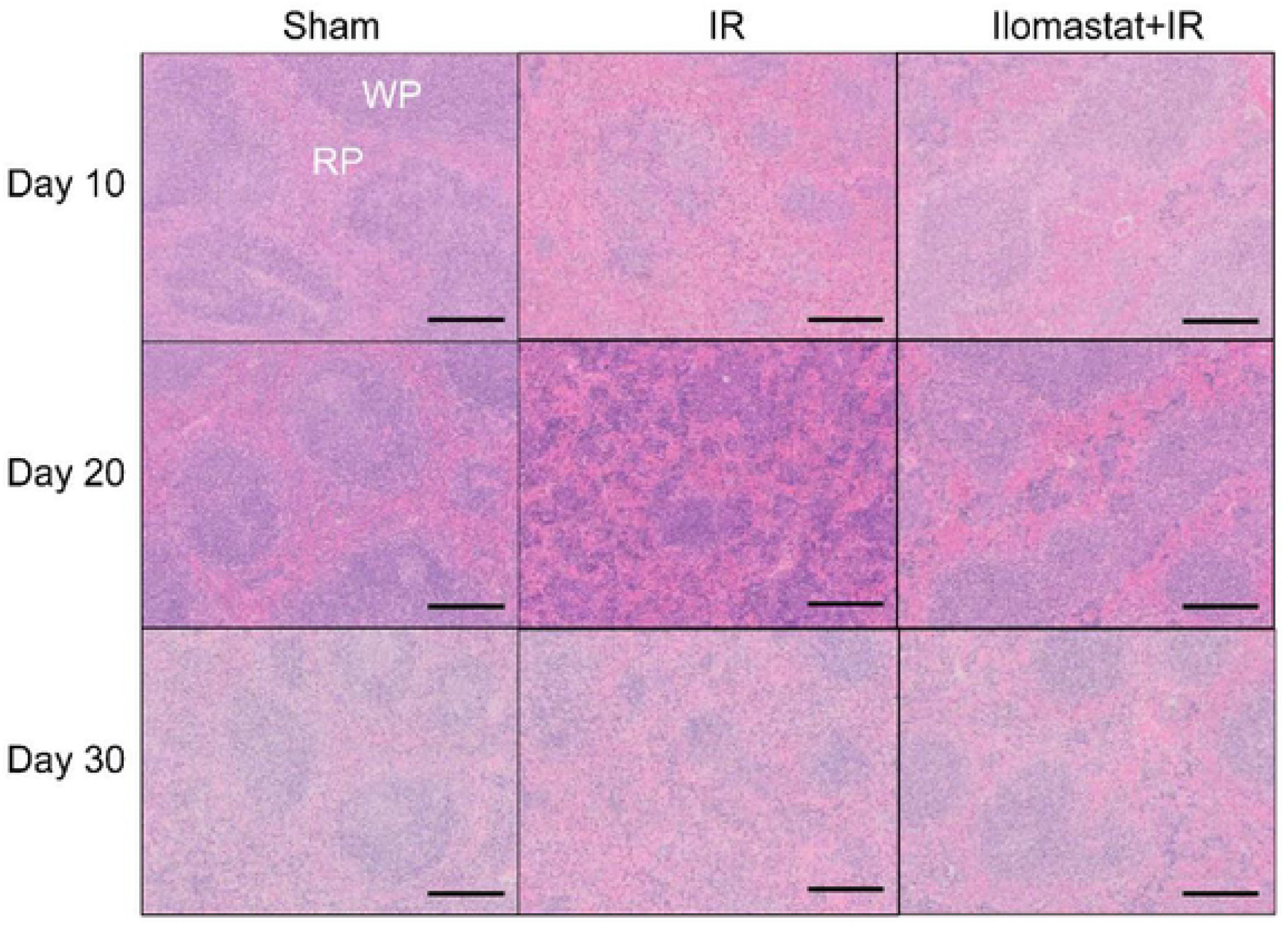
Ilomastat pretreatment improves the spleen’s pathology of irradiated mice. Mice were administrated with/without 10 mg/kg Ilomastat 2 h before exposure to 6 Gy γ-ray irradiation. H&E staining in the spleen of irradiated or unirradiated mice at day 10, 20 and 30 and the representative images were presented. Bar = 100 μm.

**Figure 6.**
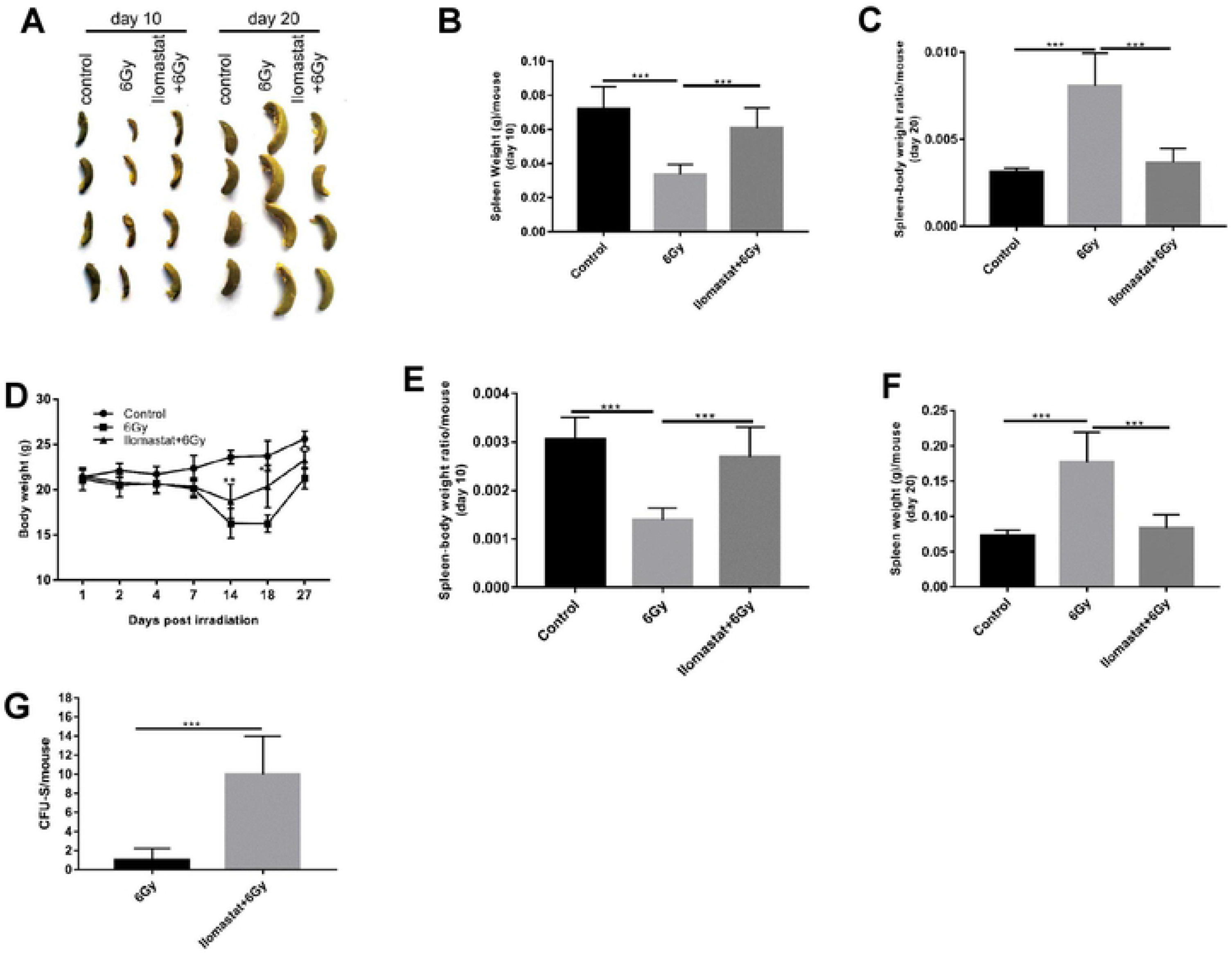
Ilomastat improves the spleen index and stimulates the splenic progenitor cells. Mice were administrated with/without 10 mg/kg Ilomastat 2 h before exposure to 6 Gy γ-ray irradiation. **(A)** The morphology of spleen extracted from mice in each experimental group. **(B-C)** The spleen weight of mice in each experimental group at day 10 and 20 after irradiation. **(D)** The body weight of mice in each experimental group at indicated days after irradiation. **(E-F)** Spleen-body weight ratio of mice in each group at day 10 and 20 after irradiation. **(G)** Spleen was fixed in Bouin’s solution to enumerate the CFUs in IR and Ilomastat + IR.All error bars indicate SEM (n = 8). \**P* < 0.05; \*\**P* < 0.01; \*\*\**P* < 0.001.

The numbers of endogenous colony forming units (e-CFUs) serve as an indicator for hematopoiesis, a crucial and indispensable factor dictating hematopoietic recovery post radiation [5, 24]. To determine the effects of Ilomastat on hematopoiesis and recovery following IR, endogenous spleen colony forming assay was performed. The number of endogenous CFUs in the irradiated mice alone was 1.0 ± 0.42, which was lower by nearly 10 folds than that in the mice pretreated with 10 mg/kg Ilomastat (10.0 ± 1.40, Fig. 6G), suggesting that Ilomastat pretreatment significantly accentuated the number of splenic colonies in the irradiated animals and stimulation of endogenous CFU-S by Ilomastat pretreatment contribute to the protection of hematopoietic injury.

## 4 DISCUSSION

The side effects of IR in patients receiving radiotherapy and the risks of exposure to IR accidentally for nuclear workers are inevitable. The methods to protect the radiation induced injuries in human are still not enough. Depletion of peripheral blood cells and immunosuppression are the main characters of ARS induced by high dose IR, which results in the death of the most irradiated animal. It is a better choice to protect victims from IR via accelerating the recovery of radiation-induced hematopietic and immunological system damages.

TGF-β1 and TNF-α are kinds of HIF. Expression and secretion of TNF-α and TGF-β1 by lymphocytes after radiation exposure was shown to be significantly increased [25]. Our present study showed that Ilomastat could significantly reduce the level of TGF-β1 and TNF-α, and increase the levels of SCF in serum. These lines of evidence suggest that MMPs inhibitor, Ilomastat, pretreatment promotes the hematopoiesis probably through stimulating the expression of HGF (SCF) and inhibiting the expression of HIF (TGF-β1 and TNF-α) in irradiated mice. A remarkable increase in CFUs and reduction of splenic atrophy at day 10 post TBI (6 Gy) with Ilomastat pretreatment also provides the evidence for protection of TBI-induced damage to the splenic compartment. In addition, the potential of Ilomastat to completely restores the cellular integrity of the BM (Fig. 2). Thereby, Ilomastat activated hematopoiesis may also facilitates the stem cell regeneration in the BM.

The majority of cells in bone marrow after receiving radiotherapy may arrest in G1 phase, resulting in a decrease of proliferation index. Results from our study demonstrating the significant decrease of BMNC in irradiated mice possibly result from an inhibition of the cell proliferation process, which is in agreement with the previous literatures [26, 27]. However, the more detailed molecular mechanism associated with cell cycle influenced by Ilomastat remains to be elucidated.

Ilomastat possesses the capacity of alleviating the loss of body weight and the atrophy of immune organs induced by radiation. Our results showed that Ilomastat had a tendency to normalize the spleen index, which implies that Ilomastat also has protection effect on the immune conditions against the side effects caused by radiotherapy and may plays a potential role in promoting hematopoiesis outside bone marrow. We also observed that the irradiated mice exhibited splenomegaly and that Ilomastat mitigated the radiation-induced splenomegaly. Splenomegaly is usually associated with disease processes that involve the destruction of abnormal RBC in the spleen. Splenomegaly may not only be caused by removal of RBC after irradiation but a possible overproduction of splenic T cells, which might also contribute to this effect. Further studies to elucidate the mechanisms of radiation-induced splenomegaly will be required.

A number of compounds acting on hematopoietic system have been reported to reduce radiation-induced injury [6]. However, the prolonged administrations of these hematopoietic growth factors and cytokines have possible concern for exhaustion of the hematopoietic stem and progenitor cells (HSPCs) reservoir, in addition to proinflammatory and immunogenic effects [28–30]. At present, except for several cytokines, for example G-CSF (FDA approved for the treatment of accidental radiation induced hematopoietic syndrome), no more drugs are been approved for application for radioprotection to hematopoietic injury in human by the FDA [4]. A series of our current experiments provided the theoretical basis to use Ilomastat as the hematopoietic activator to protect human from IR in the future.

In conclusion, our present study clearly demonstrated that Ilomastat pretreatment can effectively restore the BM cellularity and mitigated radiation-induced hematopoietic injury via decreasing the HIF and increasing the HGF, directly or indirectly, through inhibition of MMPs activation induced by IR (Illustrated in figure 7). Our results suggest that potential of Ilomastat as an alternative drug of protection during the radiotherapy or radiation accident.

**Figure 7.**
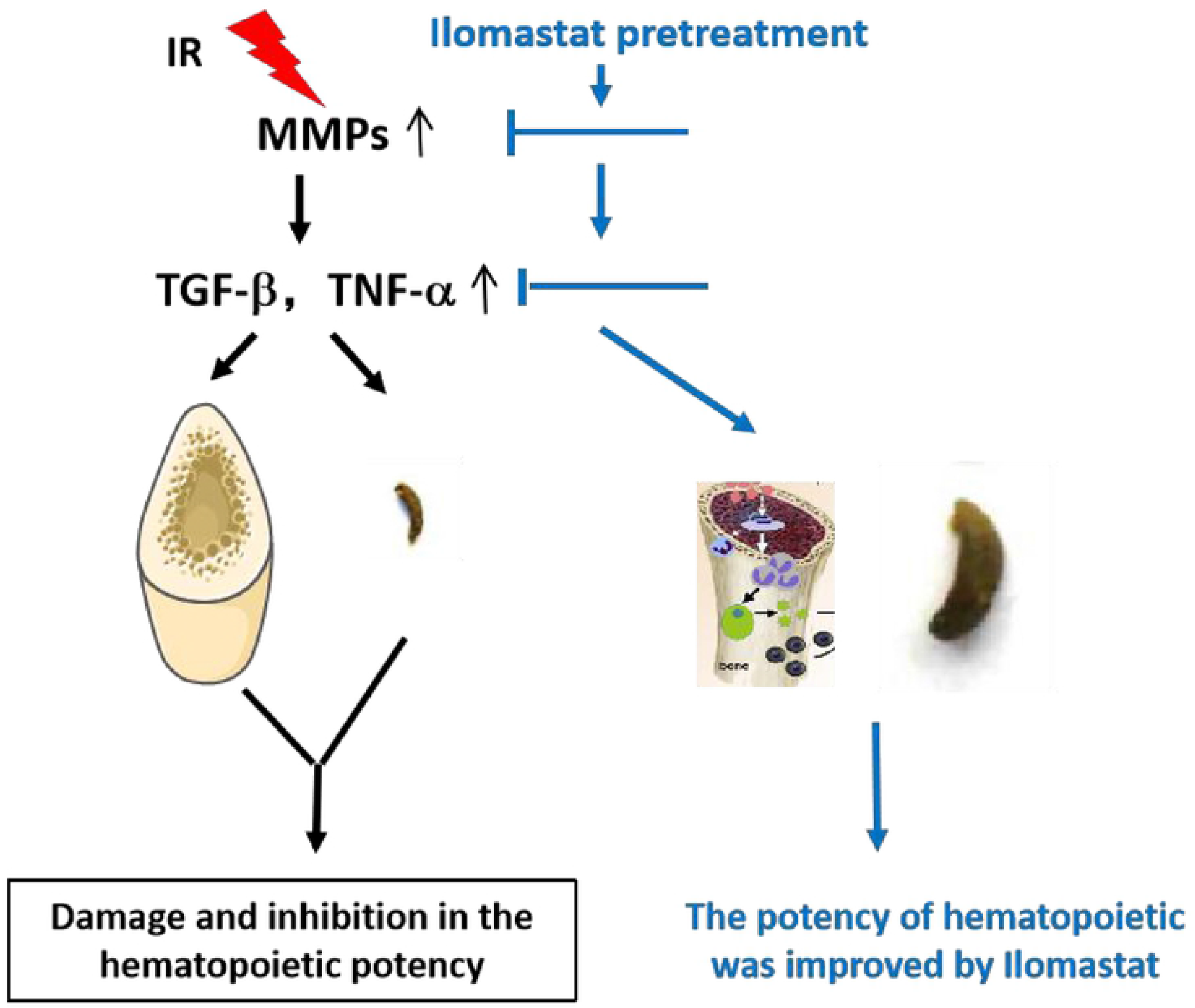
The speculated model of Ilomastat promoting the recovery of radiation-induced hematopoietic injury in mice. Ilomastat pretreatment inhibits the activation of MMPs induced by IR, which decreases the HIF and increases the HGF, directly or indirectly. Eventually, promote the recovery of radiation-induced hematopoietic injury.

## Abbreviation

MMPs: matrix metalloproteinases
IR: ionizing irradiation
TBI: total body irradiation
BM: bone marrow
BMNC: bone marrow nucleated cells
BM-MNC: bone marrow mononuclear cells
HPCs: hematopoietic progenitor cells
HSCs: hematopoietic stem cells
HSPCs: hematopoietic stem/progenitor cells
ARS: acute radiation syndrome
ECM: extracellular matrix
BMEC: bone marrow endothelial cells
TGF-β1: transforming growth factor-β1
SCF: stem cell factor
TNF-α: tumor necrosis factor-α
ELISA: Enzyme-Linked Immunosorbent Assay
e-CFU-S: Endogenous spleen colony forming unit
HIF: hematopoietic inhibitory factor
HGF: hematopoietic growth factor

## Acknowledgements

This work was supported by the Military Medical Scientific and Technological Project [AWS17J009, BWS16J007, 2018ZX09J18102], National Natural Science Foundation of China [11635013].

## Authors’ contributions

Baoquan Zhao took charge of the experimental design, data analysis and manuscript revise. Xiaoman Li performed experiments, analyzed the results and wrote the manuscript. Xingzhou Li, Dongqin Quan and Fang Zhang helped with experiments, data interpretation and manuscript preparation. Burong Hu helped with the experimental design, data analysis and manuscript revise. All authors read and approved the final manuscript.

